# Highly potent antiviral drug candidates targeting SARS-CoV-2 nsp3 and nsp5

**DOI:** 10.1101/2025.04.28.650947

**Authors:** Mathilde Galais, Guillaume Herlem, Sandy Haidar Ahmad, Sébastien Pasquereau, Ranim El Baba, Maxime Bellefroid, Estelle Plant, Stéphanie Morot-Bizot, Fabien Picaud, Georges Herbein, Carine Van Lint

**Affiliations:** Service of Molecular Virology, Department of Molecular Biology, Université Libre de Bruxelles (ULB), Gosselies, 6041, Belgium; Nanomédecine, Imagerie, Thérapeutique, EA4662, University of Franche-Comté, Besançon, France; Pathogens & Inflammation/EPILAB Laboratory, EA 4266, University of Franche-Comté, Besançon, France; Apex Biosolutions, 3 Rue de la Terre Rouge, 25220 Roche-lez-Beaupré, France; Department of Virology, CHU Besançon, Besançon, France

**Keywords:** coronavirus, SARS-CoV-2, COVID-19, HCoV-229E, Mac1, nsp3, nsp5, drug design

## Abstract

Faced with the emergence of three epidemics linked to a virus from the *coronaviridæ* family over the last 20 years (SARS-CoV in China in 2002, MERS-CoV in Arabia in 2012, and SARS-CoV-2 worldwide in 2019), the identification of new antiviral treatments is of major public health concern. As part of the development of new drugs, several molecular modeling tools such as docking, virtual screening and molecular dynamics were combined with databases to decipher potential inhibitors of coronavirus targets. We performed a structure-based design of antiviral drugs targeting SARS-CoV-2 non-structural proteins 3 and 5 (nsp3 and nsp5) based on a high-throughput virtual screening of the ZINC15 database tranches for ligand docking followed by click chemistry with a particular attention paid to relaxed structures mimicking potential in vivo interactions. Based on the above *in silico* approaches, we selected two small molecules with high affinity binding for nsp3 and nsp5 named Amb929 and Amb701, respectively. We then assessed the antiviral activity of both the Amb929 and Amb701 against SARS-CoV-2 infection taking into account their potential cytotoxicity. Although both compounds inhibited SARS-CoV-2 replication *in vitro*, Amb929 displayed the most efficient anti-SARS-CoV-2 activity. Among these two drugs, Amb929 was less cytotoxic compared to Amb701, demonstrating an optimal anti-SARS-CoV-2 response with a high selectivity index in cell culture and a high inhibition of SARS-CoV-2 replication compared to untreated condition on a model of human airway epithelium (HAE), paving the way for the development of drugs with very potent anti-coronavirus activity.

## Introduction

Human coronaviruses (HCoVs) are positive-stranded RNA pathogens that encompass seven different strains divided into two genera: alphacoronavirus (alphaCoV) such as HCoV-229E and betacoronavirus (betaCoV) such as SARS-CoV-2 (1–3). Coronaviruses are spherical enveloped viruses with a diameter around 100 nm. Briefly, they consist of a capsid formed by the nucleoprotein N (∼50 kDa) surrounded by a lipid bilayer derived from the host cell and the RNA genome inside. Three structural proteins are embedded on their surface : the membrane protein M (∼27 kDa), the envelope protein E (∼10 kDa) and the protein S (∼190 kDa), also called spike which allows the binding of the virus to its cellular receptor, human aminopeptidase N (hAPN) (4) and angiotensin-converting enzyme 2 (ACE2) (5,6) for HCoV-229E and SARS-CoV-2, respectively. Unlike the protein S which is highly variable, non-structural proteins (nsp), notably nsp3 and nsp5 of HCoVs and SARS-CoV-2, are quite stable with highly conserved regions (7). Additionally, nsp3 (PLpro, papain-like protease) and nsp5 (Mpro, main protease or 3CLpro, 3C-like protease) are viral proteases responsible for the majority of cleavages of the replicase polyproteins pp1a and pp1ab. Nsp3 is composed of eight domains (the ubiquitin-like domain 1 (Ubl1), the Glu-rich acidic domain (called “hypervariable region”), a macrodomain (named “MAC1” or “X domain”), the ubiquitin-like domain 2 (Ubl2), the papain-like protease 2 (PL2pro), the nsp3 ectodomain (3Ecto, called “zinc-finger domain”), as well as the domains Y1 and CoV-Y and two transmembrane regions, TM1 and TM2 (8–13). The highly conserved Mac1 macrodomain of nsp3 plays an essential role in the pathogenesis of coronaviruses, including HCoV-229E and SARS-CoV-2 (14). The CoV nsp5 protease processes nsp at 11 cleavage sites and is essential for virus replication. CoV nsp5 has a conserved 3-domain structure and catalytic residues (15). As both nsp3 and nsp5 play an essential role during the processing of the polyproteins they were proposed to be a major target for the development of anti-coronavirus drugs (16), a strategy that has already proved efficient for other RNA viruses such as HIV and HCV (17).

Furthermore, targeting conserved regions in nsp3 and nsp5 across the coronavirus family, could allow for the development of new antiviral drugs that could resist the high rate of mutations observed in coronavirus strains (18). The main benefit of drugs targeting nsp3 or nsp5 proteins over spike protein is that since both nsp3 and nsp5 are highly conserved and less prone to mutations their targeting might allow for a sensitivity to antiviral treatment available for several coronavirus strains from the less pathogenic alpha coronavirus HCoV-229E to the highly pathogenic betacoronavirus SARS-CoV-2. As there are currently a limited number of antiviral drugs available to treat COVID-19 infection, the search for reliable inhibitors used clinically as a potential treatment option is of paramount importance (19,20).

In the present study, we identified novel inhibitors of SARS-CoV-2 nsp3 and nsp5 by structure-based high throughput virtual screening, *in silico* click-chemistry and molecular dynamics simulations. We uncovered two small molecules with high binding affinity to nsp3 and nsp5 respectively, namely Amb929 and Amb701. We assessed the anti-SARS-CoV-2 activity of both molecules *in vitro* and as well as *ex vivo* on human airways epithelial cultures.

## Results

### Identification of new drug-like ligands docking SARS-CoV-2 nsp3 and nsp5 with high affinity

SARS-CoV-2 nsp3 is the largest SARS-CoV-2 protein and constitutes an essential component of the replication/transcription complex. We focused our structure-based design of antiviral drugs model on the conserved nsp3 Mac1 domain 6W02 (Fig 1A). The domain of SARS-CoV-2 nsp5 protease analyzed, namely 6LU7 (306 residues) was determined by X-ray diffraction at the resolution of 2.16 Å (21,22) (Fig 1B). Several significant results emerged from docking analysis and VS calculations applied to viral nsp3 (6W02) and nsp5 (6LU7). The main one was the appearance of new cavities in the solvated state which were not present in the crystal structure (Fig 1C-D). For instance, the nsp3 crystal structure 6W02 co-crystallized with the ligand ADP-ribose showed only one identified cavity, in a cleft at the top of the central β sheet (β7−β6−β3−β5). Indeed that structure shows the best score (Fig 1C,a) after VS is bound to this binding site in agreement with the unique cavity found by the Prank tool (Fig 1D,a), with a pocket score of 22.7 (27 amino acids involved). But once solvated, nsp3 showed three cavities with pocket scores of 8.9 (17 amino acids involved), 1.9 (7 amino acids), and 1.7 (7 amino acids). If top list docked ligands remain in the cleft at the top of the central β sheet (Fig 1C,b), new possibilities have also been offered in agreement with Prank tool (Fig 1D,b) (23). It follows that the protein dynamics during solvation reveal the formation of new unsuspected pockets in the crystalline state, in addition to the pocket deformations (24). This trend was confirmed with the tested nsp5 structure. The crystal structure nsp5 6LU7 (Fig 1C,c) had four calculated cavities (pocket scores 10.5; 2.1; 2.0 and 1.9, Fig 1D, c) and six when solvated (Fig 1C,c D,d). We screened a database of 205 034 compounds called the KKAA tranche of ZINC15 (25). All ligands that had the highest score from the KKAA tranches of ZINC15 and were docked in the identified cavity of nsp3 and nsp5 had rich chemistry structures. Among the top scoring output compounds, the highest scores of 13.5 (-13.5 kcal/mol) for nsp3 6W02 (ZINC000002782982) and of 11.4 (-11.4 kcal/mol) for nsp5 6LU7 (ZINC000004527915) allowed to define drug-like ligands with high binding affinity (Fig 2A).

**Fig 1.**
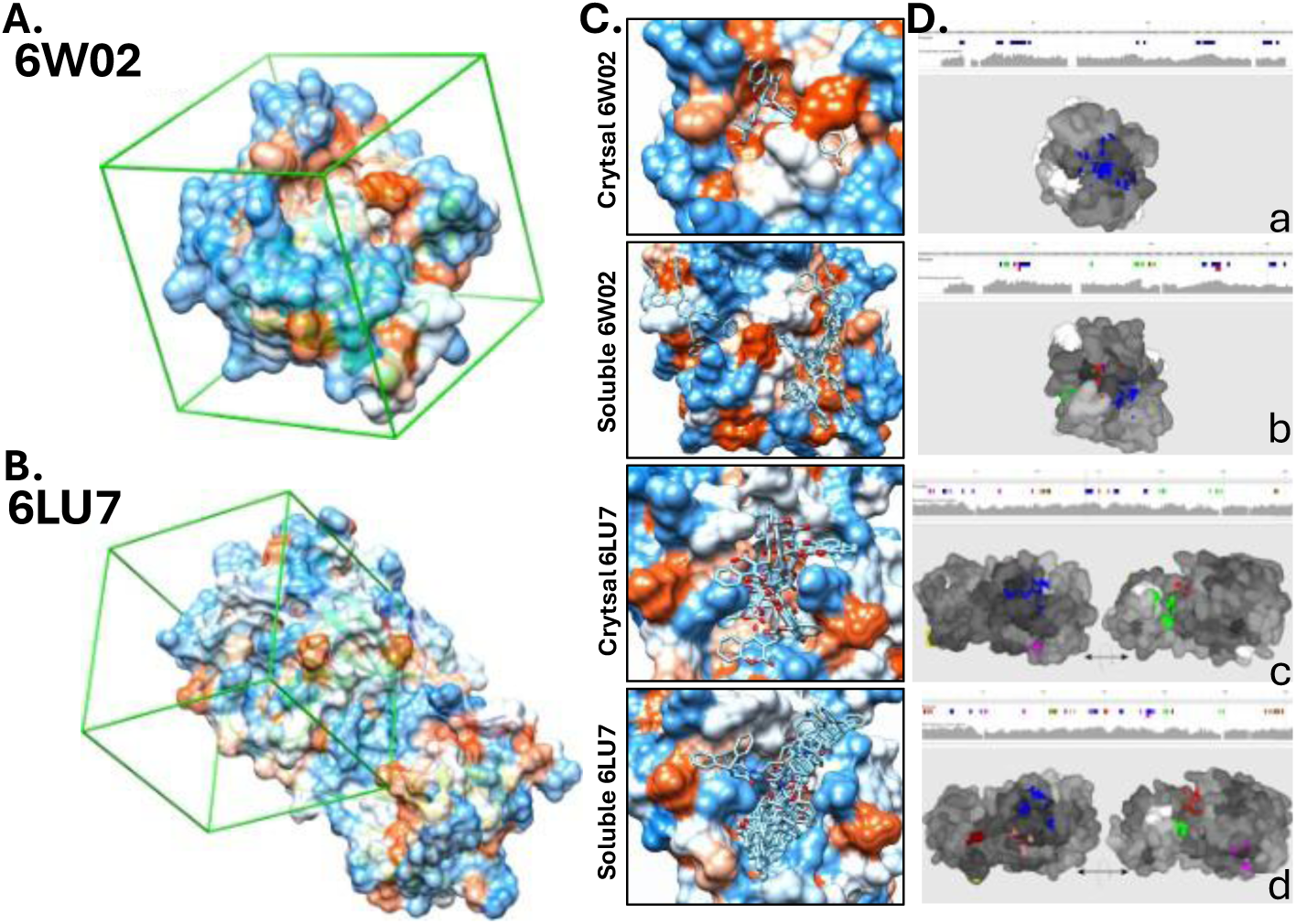
Identification of cavities from crystal and soluble structures of nsp3 and nsp5. Binding sites of SARS-CoV-2 Nsp3 (6W02) **(A)** and Nsp5 6LU7 **(B)** were predicted by P2Rank. The same box size 35 Å was used for docking all proteases (green box). In each picture, the cavity is behind the front edge of the box. Both crystal and soluble in physiological solutions structures are presented for each viral domain **(C)** with calculated corresponding binding cavities available for ligands **(D).**

**Fig 2.**
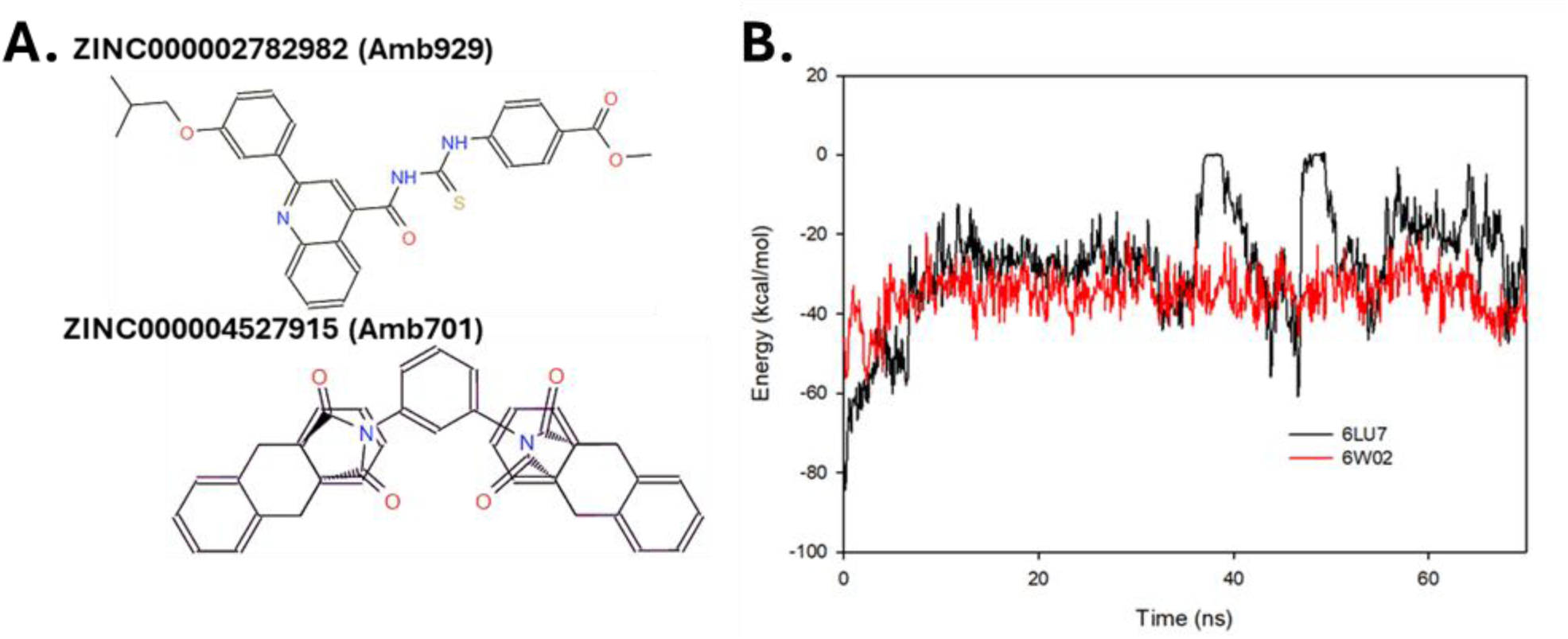
Ligands screened from KKAA tranches of ZINC15 docking with highest affinity in nsp3 and nsp5 soluble cavities. **A.** Chemical structure of best drug-like ligands docking in 6W02 and 6LU7 namely ZINC000002782982 (Amb929) and ZINC000004527915 (Amb701) respectively. **B.** Stability of the different complexes composed of new drug-like ligands docking SARS-CoV-2 nsp3 (6W02) and nsp5 (6LU7) with high affinity. Pair interaction energy (in kcal/mol) as a function of simulation time (in ns) for each best-scoring result.

To assess the stability of the different complexes composed of new drug-like ligands docking nsp3 (6W02) and nsp5 (6LU7) with high affinity we performed molecular dynamics (MD) simulations. The stability of the different complexes obtained by VS was tested under physiological conditions. For this, we placed in a simulation box containing water charged with NaCl (0.15 mol/L) each system presenting the best scoring results obtained in our studies, and at 310K thanks to molecular dynamics simulations. Once reached, the simulations were run for 70 ns to verify the degree of interaction between each ligand and its SARS-CoV-2 protein. We plot in Figure 2B the pair interaction energies obtained for each complex during this production phase. As depicted in Figure 2B, each complex exhibits a stable pair interaction (negative energy) that did not vary much during the production time. These pair interactions fluctuated from -80 to -20 kcal/mol depending on which protein were far from thermal agitation at 310 K (around 0.62 kcal/mol). The complexes determined here were thus very stable in normal conditions.

### In vitro inhibition of SARS-CoV-2 replication in cultures by new drug-like ligands docking nsp3 (Amb909) and nsp5 (Amb701) with high affinity

To assess the antiviral potential of new drug-like ligands docking nsp3 (Amb929) and nsp5 (Amb701) we performed SARS-CoV-2 inhibition tests. Treatment of VeroE6 cells with nsp3 ligand Amb929 and nsp5 ligand Amb701 showed limited and moderate cytotoxicity, respectively, with a calculated CC50 of 281 µM and 97µM respectively (Fig 3, Table 1). These results were consistent with observations of the cell culture as shown in SupplData1. Indeed, treatment with Amb929 showed little effect on VeroE6 cells at each drug concentration tested. However, VeroE6 cells treated with Amb701 revealed the occurrence of small crystalline debris in the cell culture increasing with drug concentration. Next, we tested the ability of both drugs Amb929 and Amb701 at various concentrations to inhibit SARS-CoV2-mNG replication in VeroE6 cells (MOI 0.01) during 48h of treatment. We showed by RTqPCR a significant inhibition of SARS-CoV-2 infection from 0 to 99% when increasing the concentration of Amb929 from 0 to 200 µM, resulting in an EC50 of 34.7 µM (Fig 4A, Table 1). We also observed that Amb929 inhibited SARS-CoV-2 replication by 3 logs at 200µM (Fig 4B).

**Fig 3.**
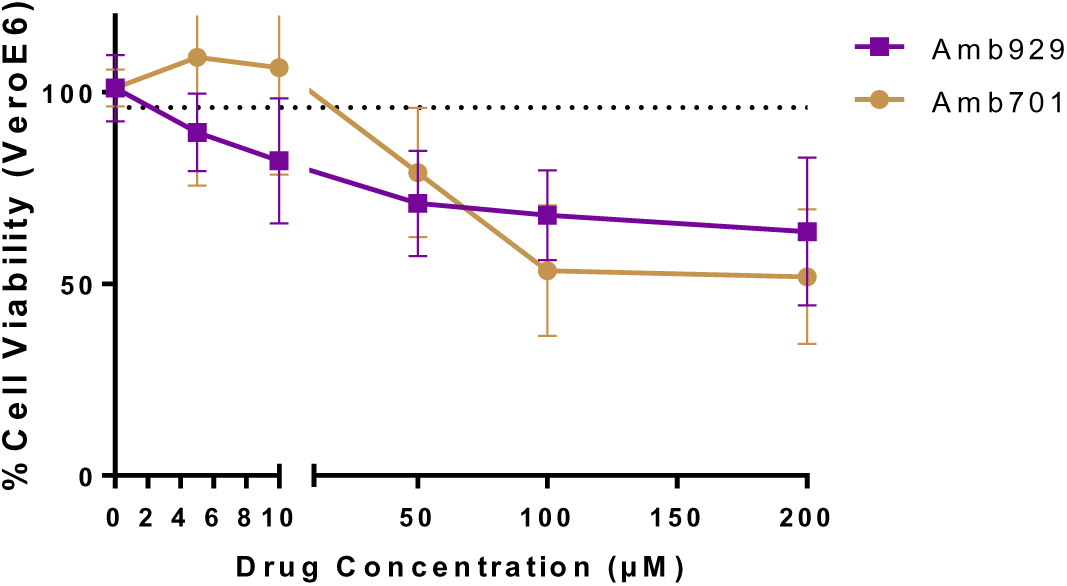
Viability of Vero E6 cells treated with Amb929 or Amb701 drugs measured by WST-1 assay. WST-1 viability assays were performed on VeroE6 cells after 48 hours of treatment with several doses of Amb929 or Amb701 (5, 10, 50, 100, 200µM). Untreated cells were used for normalization of the WST-1 viability assay. The dotted line indicates cells treated with the highest dose of DMSO (0.02%). Results are represented as mean ± SD of at least three biological replicates.

**Fig 4.**
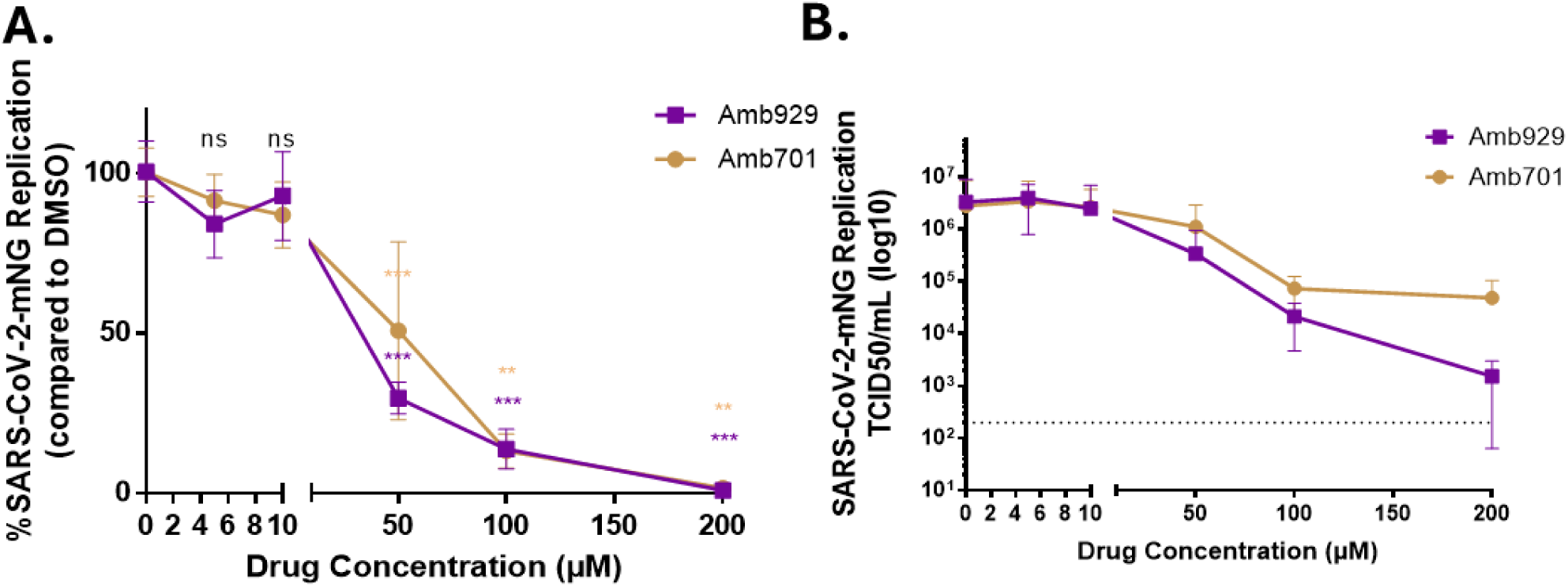
Inhibition of SARS-CoV-2-mNG replication in VeroE6 cells treated with Amb929 and Amb701 in vitro. **A.** Inhibition of SARS-CoV-2-mNG replication in VeroE6 cells treated with Amb929 or Amb701 as measured by RT-qPCR. Cells were treated by compounds at the time of infection with SARS-CoV2-mNG (MOI=0.01) and RT-qPCR were performed 48 hours postinfection. Cells treated with DMSO 0.02% were used for normalization of the assay. Results are represented as mean ± SD of at least two independent experiments done in triplicates. **B.** Inhibition of SARS-CoV-2-mNG replication in VeroE6 cells treated with Amb929 and Amb701 as measured by endpoint dilution (TCID50/mL). Endpoint dilution assays were performed 48 hours post-infection. TCID50 results are represented as mean ± SD of three independent experiments done in triplicates. The Mann-Whitney test was used as a statistical test with P ≤ 0.05 = *; P ≤ 0.01 = **; P≤0.001=***.

**Table 1.** CC50, EC50, and SI of Amb929 and Amb701 on SARS-CoV-2 replication in Vero E6 cells.

| Compound | CC50 (μM) | EC50 (μM) | SI |
| --- | --- | --- | --- |
| Amb929 | 281.9 | 34.7 | 8.12 |
| Amb701 | 97.1 | 55.4 | 1.75 |

Similarly, but to a lesser extent, Amb701 inhibited SARS-CoV-2 replication by 2 logs at 200 µM (Fig 4B). In accordance, the percentage of inhibition of SARS-CoV-2 ranges from 0 to 99% when increasing the concentration of Amb701 from 0 to 200 µM with an EC50 of 55.4 µM (Fig 4A, Table 1). However, these last results need to be taken into account with caution as we observed moderate toxicity of Amb701 molecule. We completed our observation of cell culture after treatment with either molecule by fluorescence microscopy indicating a high inhibitory effect of Amb929 on SARS-CoV-2, especially at concentrations of 50 to 200 µM (SupplData1). We calculated the selectivity index (SI) for the inhibition of SARSCoV2-mNG in VeroE6 cells for both drugs tested. Amb929 revealed a high SI of 8.12 (Table 1). In contrast, a low SI of 1.75 was obtained for Amb701 due to the moderate toxicity of this molecule observed on VeroE6 cells. Together, these results indicated that the two molecules Amb929 and Amb701 were capable of inhibiting SARS-CoV-2, with a more promising effect observed with the nsp3 inhibitor Amb929 than with the nsp5 inhibitor Amb701.

We further tested various treatment combinations with the nsp3 ligand Amb929 and the nsp5 ligand Amb701. We decided to focus on three combinations of both drugs, either a high dose of nsp3 ligand combined with a lower dose of nsp5 ligand (200µM Amb929 – 5µM Amb701) or the opposite (5µM Amb929 – 200µM Amb701), as well as a moderate dosage of both inhibitors (50µM Amb929 – 50µM Amb701) (Fig 5A). These conditions were compared to cells treated with an equivalent percentage of DMSO (0.02%) present in the culture medium. The first combination (200µM Amb929 – 5µM Amb701) seemed respectful of cellular integrity as observed by cell culture microscopy (SuplData2). The second condition of a moderate combination of both inhibitors (50µM Amb929 – 50µM Amb701) revealed an increase in the presence of precipitates in the cell culture. The latter combination of inhibitors (5µM Amb929 – 200µM Amb701) showed a lot of precipitates in the cell culture and also a fewer number of cells (SuplData2). Toxicity results indicate a moderate effect of each combination, around 50%, compared to DMSO-treated cells (Fig 5A). The level of inhibition of SARSCoV2-mNG replication in VeroE6 cells was also evaluated in the culture supernatant by RTqPCR (Fig 5B). Again, the results were compared to a treatment of 0.02% DMSO. First, the drug combination (200µM Amb929 – 5µM Amb701) presented similar levels of inhibition as single molecule Amb929 (99%). The second drug combination (50µM Amb929 – 50µM Amb701) revealed slightly better replication inhibition (80%) than the molecules used individually at this concentration (65% and 70% respectively). However, no synergic inhibition was observed for this condition. Finally, the final combination of molecules (5µM Amb929 – 200µM Amb701) showed a mean inhibition of 88% of viral replication which did not surpass inhibition levels of the molecule Amb701 at 200µM (99%). Nevertheless, the latter treatment affects a number of cells available for viral replication, as previously observed, and should be observed with caution since this treatment requires further testing as mentioned. We completed our observation of cell culture after combined treatment by microscopy indicating a high inhibitory effect of a high dose of nsp3 ligand combined with a lower dose of nsp5 ligand (200µM Amb929 – 5µM Amb701) compared to the opposite (5µM Amb929 – 200µM Amb701) (SupplData2).

**Fig 5.**
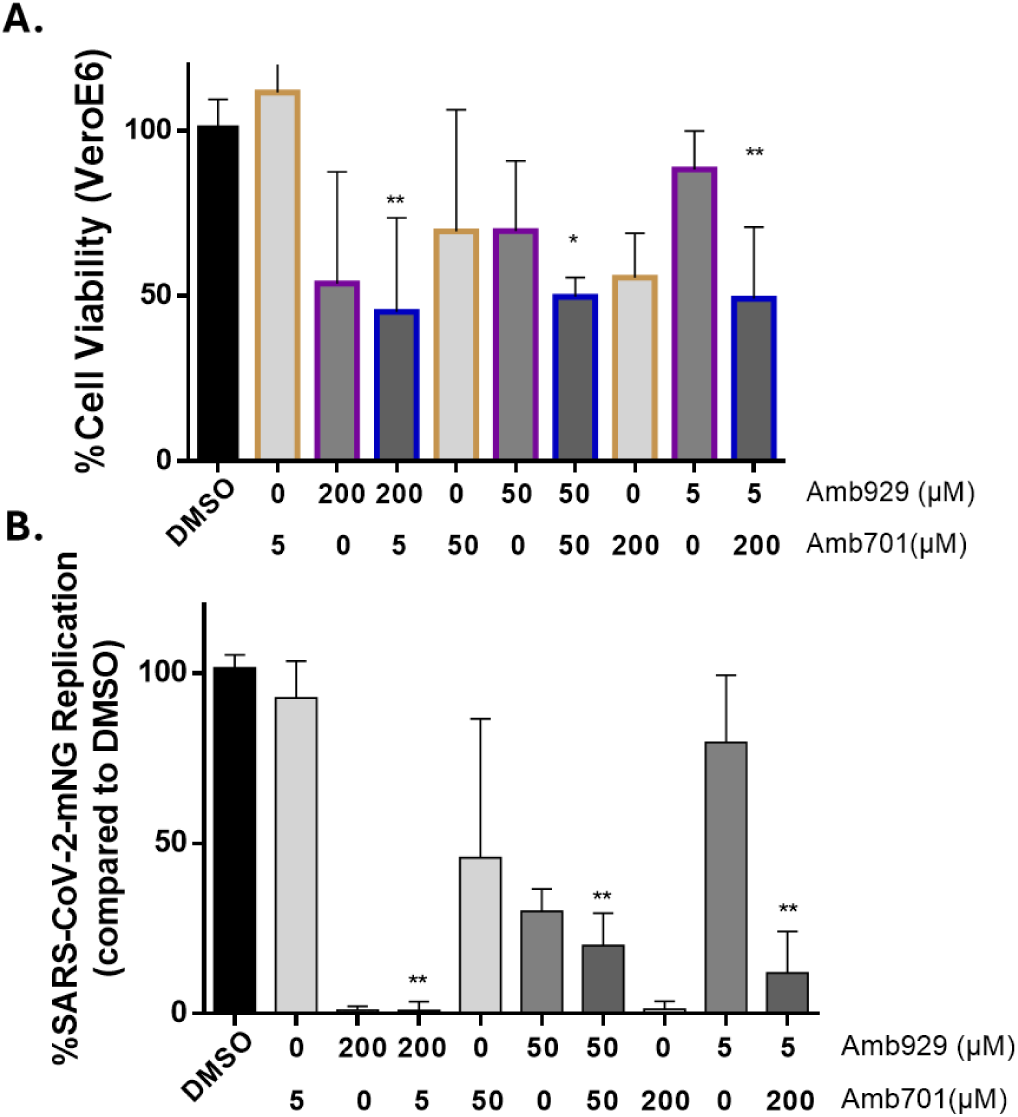
Inhibition of SARS-CoV-2-mNG replication in VeroE6 cells treated with combination of Amb929 and Amb701. **A.** Viability of Vero E6 cells treated with Amb929 and/or Amb701 drugs was measured by WST-1 assay after 48h. **B.** Inhibition of SARS-CoV-2-mNG 636 replication in VeroE6 cells treated with combination of Amb929 and Amb701 as measured by RT-qPCR. Cells were treated by a combination of compounds Amb929 and Amb701 at the time of infection with SARS-CoV2-mNG (MOI=0.01). RT-qPCR was performed on supernatant 48 hours post-infection. Control conditions were cells treated with DMSO 0.02% and used for normalization of both assays.Histogram represents mean ±SD of at least two independent experiments done in triplicates. The Mann-Whitney test was used as a statistical test with P ≤ 0.05 = *; P ≤ 0.01 = **.

### Inhibition of SARS-CoV-2 replication in a model of Human Airway Epithelium (HAE) by the new drug-like ligands docking nsp3 (Amb929) and nsp5 (Amb701)

As Amb929 and Amb701 molecules were being tested for their potential inhibitor effect of coronavirus in humans, we decided to test these molecules on a more physiological model.

Human Airway Epithelium (HAE) were infected with SARSCoV2-mNG infection (MOI 0.1) for two hours in the presence of either treatment of Amb929 or Amb701 drugs (200µM). This infection was then followed by a wash of the apical interface HAE allowing for a rapid absorption of the drug on the epithelium composed of various cell types. At 72h post treatment with Amb929 HAE showed no major alterations. In contrast after 72h of treatment with Amb701, many cellular alterations in HAE could be observed and therefore resolved our attempts to further test this drug (Suppl. Data 3). We tested the cytotoxicity of our treatment in this model via a lactate dehydrogenase activity (LDH) assay after 72h in the apical supernatant. Our results showed a moderate toxicity in HAE treated with Amb929 compared to untreated HAE, as only a 15% increase in mortality was observed (Fig 6A). After 72h, the apical supernatant was tested by RTqPCR to detect the presence of viral particles. Our results showed a 90% inhibition of replication by Amb929 treatment at 200µM after 72h compared to an untreated condition (Fig 6B). Treatment of HAE with Amb701 at 5µM, while not affecting the macroscopic state of HAE (Data not shown), did not inhibit viral replication (Fig 6B). We also tested the drug combination of 200µM Amb929 and 5µM Amb701 which revealed an interesting inhibition of SARS-CoV2 replication by 97% (Fig 6B).

**Fig 6.**
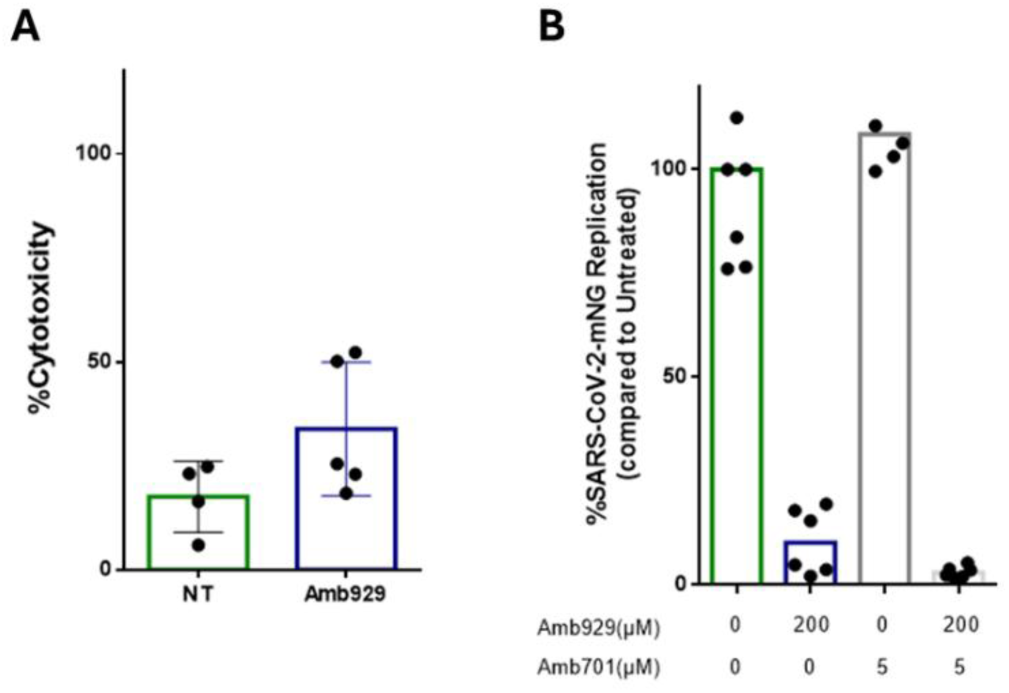
Inhibition of SARS-CoV-2-mNG replication in Human Airway Epithelium (HAE) treated with Amb929. **A** Cytotoxicity observed in apical part of HAE treated with Amb929 compared to untreated (NT) using a LDH assay. **B**. Inhibition of SARS-CoV-2-mNG replication in HAE treated with Amb929 alone or with low non-toxic concentrations of Amb701 as measured by RT-qPCR. LDH assay and RT-qPCR were performed 72 hours post-infection on apical supernatant. Untreated cells infected with SARS-CoV2-mNG were used for normalization of the assay. RT-qPCR results are represented as mean ±SD of at least four independent HAE treated or not.

## Discussion

We performed a structure-based design screening for antiviral drugs targeting SARS-CoV-2 nsp3 and nsp5 to fight COVID-19 infection. Our in silico approach was based on a high-throughput virtual screening of the ZINC15 database tranches for ligand docking followed by click chemistry with a particular attention paid to relaxed structures mimicking potential in vivo interactions. We then selected two small molecules with high affinity binding for nsp3 and nsp5 named Amb929 and Amb701, respectively. We assessed the antiviral activity of the Amb929 and Amb701 on SARS-CoV-2 infection also taking into account their potential cytotoxicity.

Although both compounds inhibited SARS-CoV-2 replication in vitro, Amb929 displayed the most efficient antiviral activity which was further confirmed on human airway epithelium cultures.

As a structure-based design of antiviral drugs targeting SARS-CoV-2 nsp3 and nsp5 was performed, we used a double strategy. First, the search of ligands with similar polarity- hydrophobicity and molecular weight collected in the ZINC15 database for docking, followed by virtual high-throughput screening calculations allowed the identification of the best candidates inhibiting nsp3 (6W02) and nsp5 (6UL7) proteins from SARS-CoV-2. Second, based on chemical fragments resulting from the best score, the ligands were carved on a measure to stick to protein cavities by click chemistry (26) with special attention paid to relaxed structures approaching living on the docking results. Indeed, the appearance of new binding cavities on the surface of nsp3 and nsp5 proteins in the physiological state, unsuspected in the crystalline state is the first evidence of the importance of our strategy. This was moderately surprising but little studied so far. In the light of our results, it therefore seemed necessary to pay rigorous attention to the solvated state of drugs in development. Current tools available for predicting protein cavities are efficient and reliable in their recent development. Indeed, there is an excellent correspondence with the docking results. The second highlight comes from the VS calculations of ligands from different tranches of the ZINC15 database. Compounds with a molecular weight between 500 and 750 Da gave the best docking scores. In our case, this concerned around 260,000 ligands, also taking into account the molecules created by the fragment growth method. However, other factors must be taken into account when performing VS calculations in the form of selection criteria such as solubility and toxicity. This greatly reduces the number of molecules but is well suited for *in vitro* tests prior to any clinical trial. Our methodology described here therefore allowed us to identify at least one high-scoring ligand that showed a broad spectrum of inhibitory action on each of the non-structural proteins nsp3 and nsp5 of SARS-CoV-2, namely Amb929 and Amb701. Moreover, it is possible to improve the score of the best-ranked ligand via *in silico* methods adding fragments by click-chemistry. However, new techniques are already emerging which use deep convolutional neural network by extending a ligand with a molecular fragment, to better match the shape of the receptor cavity (27,28).

As nsp3 and nsp5 proteases are displaying only a limited mutation rate compared with other proteins such as viral Spike, and are highly conserved through members of the coronavirus family we evaluated the broader antiviral activity Amb929 and Amb701 could have (29–31). Indeed, we assessed their antiviral activity against HCoV-229E in vitro. Both drugs inhibited HCoV-229E replication with an EC50 at 20.8µM and a 78 % reduction in viral replication for Amb929 (Suppl. Data 4B, Table 2), and Amb701 showed an EC50 of 11.9µM and 92 % reduction in viral replication (Suppl. Data 4B, Table 2). However, we measured also their toxicity on MRC5 cells. Amb929 showed the lowest cytotoxicity compared to Amb701 with a CC50 of 109.7µM and 4.4µM respectively (Suppl. Data 4A, Table 2). As previously observed in VeroE6 cells, the higher toxicity of the Amb701 molecule imposes caution on these inhibitory effects observed. Therefore, Amb929 is still a good potent inhibitor of coronaviruses infection demonstrated for both SARS-Cov-2 and HCoV-229E.

**Table 2.** : CC50, EC50 and SI of Amb929 and Amb701 on HCoV-229E replication in MRC5 cells.

| Compound | CC<sub>50</sub> (μM) | EC<sub>50</sub> (μM) | SI |
| --- | --- | --- | --- |
| Amb929 | 109.7 | 20.8 | 5.3 |
| Amb701 | 4.4 | 11.9 | 0.4 |

Moreover the inhibitory effect of Amb929 observed here was similar to that of remdesivir with an EC50 of 34.7µM and 23.1µM respectively in VeroE6 cultures (32). Other drugs targeting SARS-CoV-2 nsp3 especially its Mac1 domain have been suggested based on structure-based drug design used to identify high-affinity inhibitors (33–36). We are proposing that Amb929 besides targeting the Mac1 domain of nsp3 could also restore the host immune defense through repression of interferon production which remains to be further investigated (37–40).

Besides, as nirmatrelvir, a nsp5 protease inhibitor, is already in use to treat COVID-19 in combination with ritonavir (nirmatrelvir/ritonavir) by FDA approval (41), we are hoping the identification of Amb701 another nsp5 inhibitor will further the interest for this molecule and in its development against coronavirus infections. Indeed with newly emerging variants (variants of concern, VOCs) there is a concern that mutations in nsp5 may alter the structural and functional properties of protease and subsequently the potency of existing and potential antivirals. So far nirmatrelvir demonstrated no changes in potency for all VOCs mutants, suggesting nsp5 will remain a high-priority antiviral drug candidate as SARS-CoV-2 evolves (42). Additionally with other highly potent inhibitors of SARS-CoV-2 nsp5 and nsp3 being under development (33,43), it is our hope that Amb701 can be further improved to better match the shape of its receptor cavity.

Taken together, these results further indicated the real interest of developing new drugs targeting nsp3 and nsp5 to inhibit coronavirus infections, especially SARS-CoV-2. While further testing remains to be performed and based on the observations here presented, we consider the Amb701 drug to be too disruptive of cell integrity to be a good candidate.

While Amb929 showed only a moderate toxicity on several cell types including the human model of epithelial airways, but more importantly, it was found to efficiently inhibit SARS-CoV-2. Therefore, Amb929 which targets Mac1 domain of nsp3, appears to be the best candidate for further development of drugs with highly potent anti-coronavirus activity.

## Materials and Methods

### Structure-based design of nsp3 and nsp5 antiviral drug candidates

Our drug design method consisted of several steps: drug property prediction, molecular docking and molecular dynamics simulation.

Prior to docking and virtual screening (VS) calculations, the protein structures used and coming exclusively from experimental techniques (the worldwide protein data bank www.rcsb.org) were cleaned of ions and water molecules. Then, the binding sites were located by using the position of a co-crystallized complexing ligand, SARS-CoV-2 nsp3 6W02 (Crystal Structure of ADP ribose phosphatase of nsp3 from SARS CoV-2 in the complex with ADP ribose) (44)and sp5 6LU7 (The crystal structure of COVID-19 main protease in complex with an inhibitor N3) (21). Briefly, a machine learning method was used based on local chemical neighborhood ligand ability centered on points placed on a solvent-accessible protein surface. Points with a high ligand ability score are then clustered to form the resulting ligand binding site. These pockets are circumscribed to parallelepipedal box large enough for conformational search. Thus, any ligands from a database able to accommodate to a cavity are identified and the best conformation by stabilizing the complex achieved is found. Mainly, as implemented in QuikVina-w (a major revision of Autodock Vina), the box search space is carried out by global and local optimization algorithms, in the forms of Markov chain Monte Carlo sampling method and completed and accelerated by using local search with BFGS optimization (45,46). It is based on the score function of AutoDock Vina, the accelerated search of QVina 2, and adds an in-depth search for a large search space. It allows a search thread to communicate with other nearby threads to increase the speed and sensitivity of that search thread. This communication uses a global buffer that keeps the high-quality search history of all threads to converge more efficiently and quickly to a solution. A virtual screening (VS) is then carried out via a script passing the ligands in the box one by one, then ranking the scores obtained. The scores are given in absolute value and correspond to minus the Gibbs free energy in kcal/mol.

The method based on click-chemistry reactions promotes the growth of a ligand in the protein cavity to optimize binding affinities (47–49). The process begins by placing a chemical fragment in the pocket and adding chemical groups to increase interactions with residues of the protein cavity guided by the FOG algorithm. Structural optimizations are performed after each addition to keep the conformation with the lowest Gibbs free energy (the SMOG2016 scoring function). A hydrogen atom of the current ligand is randomly selected to attach the new fragment which is chosen according to the FOG transition probabilities of the previous fragment. Amongst three growth options, SEED mode was used in this work. It relies on the best complexing ligand previously obtained from VS to improve its score by adding fragments rather than in DENOVO mode where the growth starts from a randomly chosen fragment.

In addition to crystal structures, optimized ones upon CHARMM force field used by the NAMD code were produced to get closer to the biological conditions (50). Classical MD simulations are useful tools able to relax protein structures in reliable conditions. For this, each system was solvated in a periodic water box whose dimensions were chosen large enough to prevent protein interactions with the one directly adjacent in the surrounding periodic boxes. To work as close to biological conditions, NaCl ions were added at a normal salt concentration of 0.15M to the solvated system. To model the water molecules, the TIP3P scheme was used to reproduce the correct biological environment. All solvated proteins were gradually equilibrated to reach 310K using MD simulations (NAMD 2.12 package) and then studied during production runs of 30 ns (51). CHARMM36 force-field optimization parameters were used in all simulations (52).

During the simulations, temperature and pressure were kept constant at 310 K (Langevin dynamics) and 1 atm (Langevin piston), respectively. Long-range electrostatic forces were evaluated using the classical particle mesh Ewald (PME) method with a grid spacing of 1.2 Å, and a fourth-order spline interpolation. The integration time step was equal to 2 fs and no constraint was imposed on any atoms of the system. When no molecular force field existed (particularly for ligands obtained through chemical click), the simulation parameters were obtained by building the molecular force field with the Force Field Toolkit package of the Swissparam program (53).

### Chemical ligands targeting SARS-CoV-2 nsp3 and nsp5

Chemical ligands targeting SARS-CoV-2 nsp3 and nsp5 were purchased from Ambinter (Orléans, France): Amb1934929 (ZINC000002782982) and Amb2618701 (ZINC000004527915) to target nsp3 6W02 and nsp5 6LU7, respectively. After purification, the drugs were stored in solid form at -20°C before preparation. Drugs were diluted in DMSO at a concentration of 10 mM. The 10 mM stock was further diluted in cell culture media at final assay concentration at the time of use (5, 10, 50, 100 or 200µM).

### Viruses preparation

The infectious clone SARS-CoV-2 mNeonGreen (icSARS-CoV-2-mNG) strain was provided by Dr. Pei-Yong Shi from University of Texas Medical Branch (UTMB) (54). SARS-CoV-2- mNG stock virus (7 log TCID50/mL) was propagated in Vero E6 cells (ATCC) in DMEM media supplemented with 2% FCS. Viral titers were determined by endpoint dilution assays on Vero E6 cells and stored at -80°C until use. HCoV-229E was isolated from nasal and throat swabs collected from a man with mild upper respiratory illness (Human Coronavirus 229E ATCC VR-740) and propagated using MRC5 cells (RD Biotech, Besançon, France). HCoV- 229E stock virus (5 log PFU/mL) was prepared in MRC5 cells in DMEM media supplemented with 10% FCS.

### Viral replication inhibition assay

Vero E6 cells were treated with compounds at the time of infection by SARS-CoV-2-mNG at MOI=0.01 in DMEM supplemented with 2% FCS. The viral inoculum was removed after a 2- hour incubation and the cells were overlaid with fresh DMEM media supplemented with 10% FCS containing the diluted compounds. After a 48-hour incubation at 37 °C, supernatants were collected and frozen at -80°C prior to quantification and titration by RT-qPCR and endpoint dilution assay (TCID50/mL). RT-qPCR was performed on the collected supernatants. RNA was extracted from the supernatant using an RNA extraction kit QiAmp viral RNA kit (Qiagen ID 52906). RTqPCR was performed using TaqMan Fast Virus 1-Step MasterMix (ThermoFisher 4444432) and SARS-CoV2 probes (forward (IDT 10006888); reverse (IDT 10006890); probe (IDT 10006892). Three independent replication assays were performed in triplicates for these viral replication assays. Classic tissue culture infectious dose 50% (TCID50/mL) in Vero E6 cells was performed in triplicates on viral stocks and samples collected using the Reed & Muench statistical method.

Air-liquid cultures of primary human airway epithelial (HAE) cells of bronchial origins from healthy donors were obtained from Epithélix (MucilAir) and cultured with MucilAir medium (Epithélix). The apical side of the HAE cells was washed prior to infection by SARSCoV2- mNG (MOI 0.1) and treatment, following the manufacturer’s instructions. Virus and treatment were simultaneously applied on the apical side of the cells (diluted in 200µL of MucilAir medium). Drug treatment was added at the indicated concentration for 2h during virus infection. The apical interface HAE was washed after infection and treated with a small volume of medium-containing treatment (200µL).

To evaluate the effect of compounds *in vitro*, MRC5 cells were treated with compounds diluted in culture media at the time of infection by HCoV-229E at MOI =0.1 . The antiviral compounds were maintained with the virus inoculum during the two-hour incubation period. The inoculum was removed after incubation and the cells were overlaid with culture media containing diluted compounds. After 48 h of incubation at 37°C, supernatants were collected to quantify viral loads by plaque-forming unit (PFU) assay, as previously described (55). Briefly, MRC5 cells were infected with the supernatant for 2 hours and then overlaid with culture media containing agarose (1%). Cells were incubated for 48 hours. Plaque formation was observed under a light microscope. Cells were stained with MTT for 1 hour and plaque-forming units were counted.

### Cytotoxicity assays

The WST-1 metabolic assay was performed on Vero E6 cells treated for 48 hours with compounds, following the manufacturer’s instructions (Roche Diagnostics GmbH, Mannheim, Germany). Cytotoxicity assays were performed in duplicates. LDH assay was performed and percentage cytotoxicity calculated on apical supernatant from HAE according to the manufacturer’s instructions (Promega JS2381). The cytotoxicity of the selected compounds was assessed in MRC5 cells using an MTT assay. Briefly, cells were treated with compounds for 48 hours. The culture media was replaced and the cells were incubated with MTT (1.2mM, Life Technologies, Eugene, OR, USA) for 2 hours. Formazan crystals were dissolved by addition of acidic isopropanol and cell viability was measured by a spectrophotometer (Bio-Rad, Hercules, CA, USA).

*Effective concentrations, cytotoxic concentrations and selectivity index calculations*.

Four-parameter logistic regression (IC50 Toolkit IC50.org) was used to fit the dose-response curves and determine the 50% effective concentrations (EC50) of the compounds that inhibit viral replication by 50%. The cytotoxic concentration 50 (CC50) is the concentration of the compound that reduced cell viability by 50% and was calculated using four-parameter logistic regression (IC50 Toolkit IC50.org). This index was calculated for each compound and cell type by the same method. The selectivity index (SI) was calculated by dividing CC50 by EC50 for each compound and virus tested. The selectivity index is an indicator that measures the window between cytotoxicity and antiviral activity. A high selectivity index reflects a low toxicity at concentrations that show an effective antiviral activity. It also indicates that there is an important range of concentrations showing an effective antiviral effect with minimal toxicity. Conversely, a low selectivity index would indicate an important toxicity for concentrations with effective antiviral activity.

### Microscopy

After treatment, cells were fixed by 4% formaldehyde solution and nuclei were stained by DAPI. Triplicates of each condition was then assessed by fluorescent microscopy (Axio Observer Z1) and field acquisition. Images were then processed with Cell Profiler^TM^ cell image analysis software to determine the number of infected cells (Neon Green staining) per field compared to the number of nuclei in the field (DAPI staining).

### Statistical analysis

Mann-Whitney test was used to compare each condition to a control treatment and obtain P value of dataset. Statistical analysis were performed using GraphPad Prism version 6.01 for Windows.

## ACKNOWLEDGEMENTS

Calculations were performed at the supercomputer regional facility Mesocentre of the University of Franche-Comté with the assistance of K. Mazouzi. This work was granted access to the HPC resources of IDRIS, Jean Zay supercomputer, under the allocation 2020 - DARI AP010711661 made by GENCI. We would like to express our gratitude to the IDRIS team (S. Requena, P.-F. Lavallée, R. Lacroix and S. Van Crienkingen), which was able to be very reactive to our request in a very tense pandemic climate, without whom this work would not have been possible. This work was supported by grants from the University of Franche-Comté (UFC) (CR3300), the Région Franche-Comté (2021-Y-08292 and 2021-Y-08290) and the ANR (Agence Nationale pour la Recherche) ControlCovid project to Georges Herbein. SHA is supported by an ANR grant. REB is supported by a grant of Apex Biosolutions. CVL acknowledges funding from the Belgian National Fund for Scientific Research (F.R.S-FNRS, Belgium); the French INSERM agency “ANRS/Maladies infectieuses émergentes”; the “Télévie” program of the F.R.S.-FNRS; the “Fondation Roi Baudouin”; the Internationale Brachet Stiftung (IBS); The “Amis des Instituts Pasteur à Bruxelles”, asbl.; and the University of Brussels (ULB). C.V.L. is “Directeur de Recherches” of the F.R.S-FNRS. The laboratory of C.V.L. is part of the ULB-Cancer Research Centre (U-CRC). M.G. is a fellow of the EU Marie Skłodowska-Curie COFUND action (No 801505). M.B. is funded by a fellowship from the Belgian « Fonds pour la formation à la Recherche dans l’Industrie et dans l’Agriculture (FRIA) (F.R.S.-FNRS) ». The funders had no role in the data collection, analysis, patient recruitment, or decision to publish.

## Notes

### Competing Interest Statement

The authors have declared no competing interest.

